# Reduction of aggressive behaviour following hypothalamic deep brain stimulation: involvement of 5-HT_1A_ and testosterone

**DOI:** 10.1101/2023.03.20.533520

**Authors:** Flavia Venetucci Gouveia, Mustansir Diwan, Raquel CR Martinez, Peter Giacobbe, Nir Lipsman, Clement Hamani

## Abstract

**Background:** Aggressive behaviour (AB) may occur in patients with different neuropsychiatric disorders. Although most patients respond to conventional treatments, a small percentage continue to experience AB despite optimized pharmacological management and are considered to be treatment-refractory. For these patients, hypothalamic deep brain stimulation (pHyp-DBS) has been investigated. The hypothalamus is a key structure in the neurocircuitry of AB. An imbalance between serotonin (5-HT) and steroid hormones seems to exacerbate AB.

**Objectives:** To test whether pHyp-DBS reduces aggressive behavior in mice through mechanisms involving testosterone and 5-HT.

**Methods:** Male mice were housed with females for two weeks. These resident animals tend to become territorial and aggressive towards intruder mice placed in their cages. Residents had electrodes implanted in the pHyp. DBS was administered for 5h/day for 8 consecutive days prior to daily encounters with the intruder. After testing, blood and brain were recovered for measuring testosterone and 5-HT receptor density, respectively. In a second experiment, residents received WAY-100635 (5-HT_1A_ antagonist) or saline injections prior to pHyp-DBS. After the first 4 encounters, the injection allocation was crossed, and animals received the alternative treatment during the next 4 days.

**Results:** DBS-treated mice showed reduced AB that was correlated with testosterone levels and an increase in 5-HT1_A_ receptor density in the orbitofrontal cortex and amygdala. Pre-treatment with WAY-100635 blocked the anti-aggressive effect of pHyp-DBS.

**Conclusions:** This study shows that pHyp-DBS reduces AB in mice via changes in testosterone and 5-HT1_A_ mechanisms.

**HIGHLIGHTS:**

- Posterior hypothalamus DBS reduces aggressive behavior in mice
- Aggressive behavior was correlated with plasma testosterone levels
- DBS increased 5-HT1A receptor density in the orbitofrontal cortex and amygdala
- Pre-treatment with 5-HT1A antagonist (WAY) blocked the anti-aggressive effect of DBS

## INTRODUCTION

Aggressive behaviour is a complex behavioural syndrome characterized by excessive motor activity (e.g., pacing, restlessness), verbal (e.g., yelling, shouting), and/or physical aggression (e.g., self-injury, grabbing, hitting others, destroying property) [1,2]. These behaviours are commonly observed in children with neurodevelopmental disorders [3,4], patients with neurodegenerative diseases of ageing [5,6] and those with psychiatric disorders [7,8]. In addition to suffering, these symptoms comprise a leading cause of institutionalization, reducing the quality of life of patients and caregivers [8,9]. The treatment of aggressive behaviours largely involves pharmacotherapy targeting the dopaminergic (i.e., antipsychotics) and serotonergic (i.e., antidepressants) systems, as well as behavioural therapies [10–12]. Although most patients tend to have a satisfactory response, a small subset continues to present aggressive behaviours even after all treatment options are exhausted. These patients are considered to be treatment-refractory [13,14]. Refractoriness to treatment is especially observed in patients with severe intellectual disabilities who often cannot receive behavioural therapies [15,16]. Neuromodulation therapies, including deep brain stimulation (DBS), have been investigated as alternative treatment modalities in this selected group of refractory patients [17–24].

DBS consists of the implantation of electrodes in deep brain targets to deliver electrical current and modulate dysfunctional neurocircuitries [25–28]. Overall, DBS has been shown to induce focal and widespread changes in neuronal activity along with the development of multiple forms of plasticity [25,26,29–31]. In clinical trials including a small number of patients, DBS targeting the posterior area of the hypothalamus (pHyp) was associated with an important reduction in aggressiveness [17–23]. To date, the mechanisms through which DBS exerts its anti-aggressive effects remain largely unknown.

The hypothalamus is considered to be a key component of the autonomic and limbic systems, being responsible for controlling homeostasis and motivated behaviours, including aggressiveness [17,32]. The activation of GABAergic basal forebrain projections to the habenula decreases neuronal firing in this later structure, subsequently altering midbrain serotonergic and dopaminergic transmission [33,34]. This ultimately results in a decreased serotonergic and increased dopaminergic drive to the prefrontal cortex (PFC), enhancing an excitatory transmission from this structure to the amygdala [35–38]. The amygdala is highly functionally connected with the hypothalamus. Hyperactivity of the amygdala results in the activation of the hypothalamus-pituitary-gonadal axis, increasing the production and release of testosterone [39,40] and, consequently, aggressive behaviour [41–43]. Increased testosterone levels lead to a reduction in serotonergic availability in the limbic system, supporting that testosterone modulates the serotonergic activity and further contributes to the maintenance of aggressiveness [44–46]. In preclinical settings, the mouse model of escalated aggressive behaviour is considered a standard test for investigating the neuroanatomical basis, mechanisms and possible new therapies to treat AB [47].

We delivered posterior hypothalamus (pHyp-DBS) to mice and studied the effects of this therapy in a model of escalated aggressive behaviour, including the involvement of testosterone and serotonin (5-HT).

## METHODS AND MATERIALS

### Animals and husbandry

All procedures were approved by the Sunnybrook Research Institute Animal Care Committee and were in agreement with the guidelines of the Canadian Council on Animal Care and the Animals for Research Act of Ontario. Nine-week-old male C57BL/6 mice (Charles River, stock #027) were used as residents, and nine-week-old female BALB/c mice (Charles River, stock #028) were used as partners. Ten-week-old male BALB/c mice (Charles River, stock #028) were used as intruders. Upon arrival, animals were transferred to a housing room with an inverted light cycle (12h, lights off at 7 am) and optimal temperature and humidity conditions. Mice were then placed in regular cages containing nesting material and environmental enrichment, with free access to food and water. Each resident mouse was housed with one female. Intruders were group-housed with an average of 3 animals/cage.

### Study Design

This study involved two sequential experiments to firstly test the efficacy of pHyp-DBS and investigate potential changes in 5-HT and testosterone expression following stimulation, and secondly to test the possibility of blocking this effect with a 5-HT antagonist using a crossover design. For both experiments, females received tubal ligation surgery 3 weeks before being housed with male mice to prevent pregnancy while maintaining a normal estrous cycle. After surgical recovery, each female was housed with one male for the development of male territorial behaviour. Details of all methods used in this study may be found in the Supplementary Materials.

In the first experiment, 2 weeks after being housed with a female, males were randomly assigned to receive surgery for the implantation of electrodes targeting the pHyp or control surgery (the exact same procedures with no electrode implants; n=8). Following surgery, animals were allowed to recover for one week in their home cage with the female. Mice with electrode implants were then randomly assigned to receive either active DBS (n=9) or Sham stimulation (no current, n=8) for 5 hours per day prior to behavioural testing.

Resident mice were tested in the Resident-Intruder Test (RIT) in 8 distinct encounters with intruders, followed by Open Field Testing (OF) for the assessment of locomotor activity. At the end of the experiments, blood was collected for the analysis of plasma testosterone. Brains were recovered for the study of 5-HT receptors and the verification of electrode placement. Figure 1A illustrates the study design of the first experiment.

**Figure 1.**
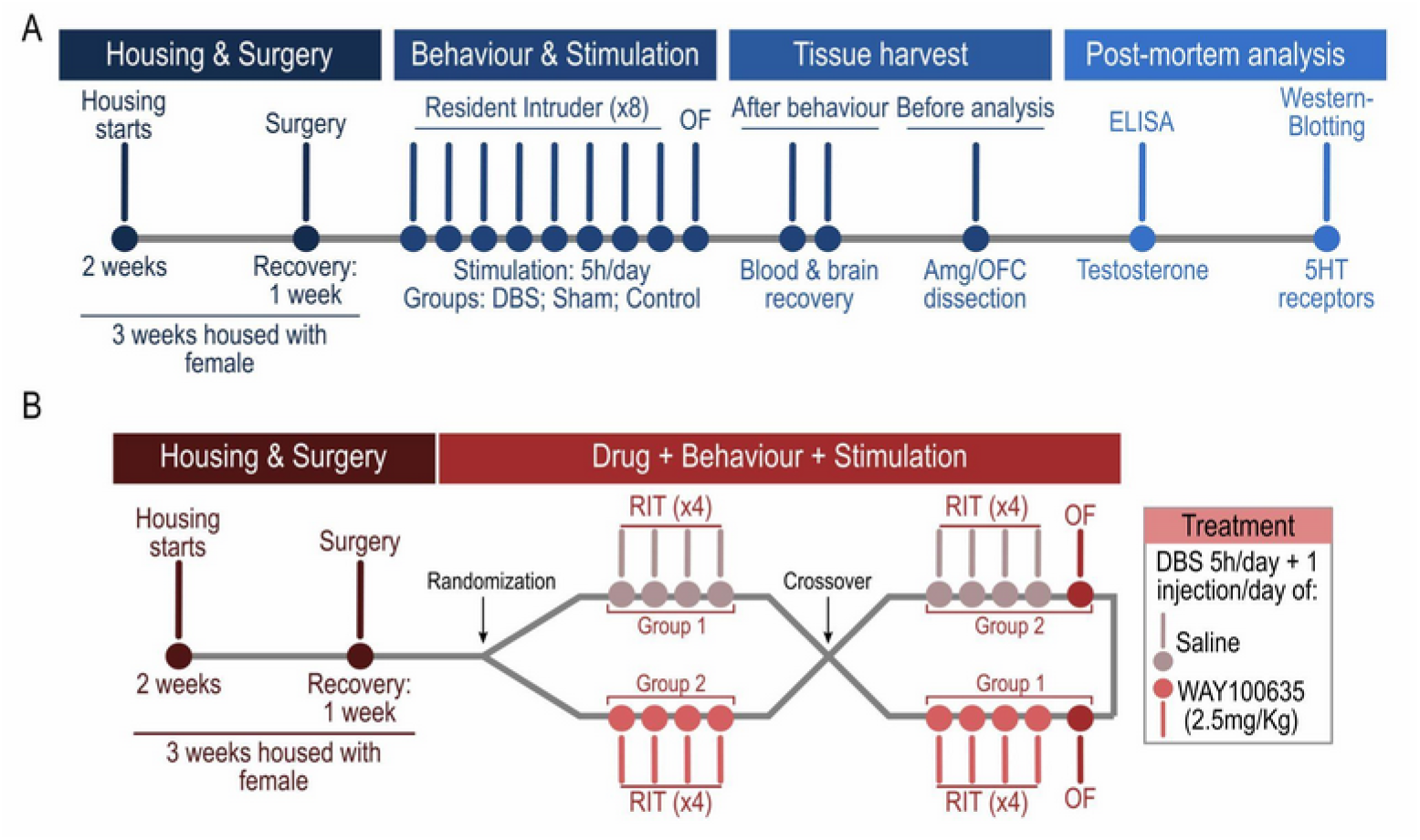
Study design. **A**. Timeline of the first experiment. **B**. Timeline of the second experiment. Abbreviations: Amg: amygdala; DBS: deep brain stimulation; ELISA: Enzyme-Linked Immunosorbent Assay; OFC: orbitofrontal cortex; OF: Open field test; RIT: Resident-intruder test.

Housing, surgical procedures and DBS administration for the second experiment were similar to the ones described above. Mice given DBS were randomly assigned to receive either the 5-HT_1A_ antagonist WAY-100635 (WAY; n=7) or saline injections (n=7) immediately prior to 5h of active stimulation. Thereafter, animals were tested in the RIT during 4 distinct encounters with intruders. The injection allocation was then crossed so that animals receiving the WAY were given saline and vice versa for another 4 encounters. Following the experiments, brains were recovered to verify electrode placement. Figure 1B illustrates the study design of the second experiment.

Following brain recovery, 15µm coronal sections covering the posterior hypothalamic area were collected and stained with cresyl-violet. The location of DBS electrode tips may be found in Supplementary Figure 1.

### Behavioural tests

Behavioral testing was conducted under red light (active time of the animals’ day/night cycle), as animals were kept in an inverted light cycle. Sessions were video-recorded and analyzed by an expert observer blinded to treatment allocation using the EthoWatcher® software [48].

#### Resident-Intruder Test

Resident mice were housed with a female for 3 weeks for the development of territorial aggression [49,50]. During the testing, residents were exposed to an intruder (i.e. unfamiliar adult mouse of the same sex) behind a protective screen for 2 min prior to the actual confrontation, which was recorded for 5 minutes. Behaviours were categorized as aggressive (i.e. bites/attacks, threats, mounting, pursuit), social (i.e. whole body social sniffing, whole body social grooming), and non-social behaviours: (i.e. self-grooming, exploring, eating, digging, resting).

#### Open Field Test

Mice were placed in the open field apparatus (50cm x 50cm x 38cm) and allowed to explore it freely for 5 minutes. The total distance (cm) travelled during the test and the average speed (cm/sec) were scored.

### Tubal Ligation

Female mice received tubal ligation surgery to prevent pregnancy 4 weeks prior to being housed with male mice. This surgery was selected over ovariectomy to preserve normal hormone cycling and facilitate mating behaviour. Under Isoflurane anesthesia and analgesia (Meloxicam, 5mg/Kg, SC), a lateral midline incision was made, and the ovary and fallopian tube were exposed. Two ligatures 0.5cm apart were tied, and the tubal area in between was resected. The abdominal cavity was closed, and the procedure was repeated on the contralateral side. Dietary supplements and analgesia were provided daily for 5 days following surgery.

### DBS surgery and stimulation

Stereotaxic surgeries were performed for electrode implant and control surgery, as previously reported [51,52]. Under Isoflurane anesthesia and analgesia (Meloxicam, 5mg/Kg, SC), one stainless steel electrode (0.5 mm lead exposure, Plastics One model 333/3) was implanted, targeting the posterior hypothalamic nucleus in the right hemisphere (ML: 0.2; AP: -2.4; DV: -4.3) [53] and 2 stainless-steel ground/anchoring screws over the parietal cortex. DBS electrodes and screws were fixed using an adhesive cement system. Control surgeries omitted electrode implants. Animals were allowed to recover for one week before behavioural testing. Electrical stimulation was performed for 5 h/day prior to behavioural testing. A connection cable (Plastics One, model 335–340/3) was used to connect the electrodes to a stimulus isolator (WPI, model A365) and the stimulator (AMPI, model Master-8). Animals in the DBS group received stimulation at 100Hz, 60μs, and 100μA, while animals in the Sham and control groups received no stimulation.

### Drug Administration

WAY-100635 maleate (2.5 mg/kg SC; Tocris) was diluted in saline and administered to the animals 30 minutes before stimulation onset on each behavioural testing day. Normal saline solution (NaCl 0.9% SC) was used as vehicle. Animals were allocated to receive either saline (group 1) or WAY (group 2) injections in the first half of the experiment. Drug allocation was crossed over during the second half of the experiments.

### Western Blot

Brains were recovered and stored at −80°C until neurochemical analysis. The orbitofrontal cortex (infra-limbic and medial-orbital cortex) and amygdala samples were bilaterally dissected. Protein extracts were separated by sodium dodecyl sulphate–polyacrylamide gel electrophoresis followed by transfer onto nitrocellulose membranes. Membranes were blocked with a buffer at 4°C overnight prior to being incubated with primary antibodies anti-5-HT_1A_ (Abcam #ab227165, rabbit pAb, 1:1000), anti-5-HT_1B_ (Abcam #ab13896, rabbit pAb, 1:1000), and anti-tubulin-β3 loading control (BioLegend #801202, mouse mAb, 1:2000). Membranes were then incubated with HRP-linked secondary antibodies (CST #7074, anti-rabbit IgG, 1:2000; or CST #7076 anti-mouse IgG 1:2000), followed by incubation with SignalFire™ ECL Reagent substrate solution, and imaged with a MicroChemi 4.2 unit (DNR Bio-Imaging Systems) using GelCapture Chemi software.

### Enzyme-Linked Immunosorbent Assay (ELISA)

Blood samples were collected in EDTA-coated capillary blood collection (Microvette® CB 300) tubes and immediately centrifuged (1000 rpm, 4°C, 10 minutes) for plasma collection and stored at −80°C until the analysis. Plasma samples, blank controls and a 6-point standard curve were added to the pre-coated microplate in duplicates. An HRP-conjugate and anti-testosterone antibody (Biomatik, #EKC40192) were added to each well, the microplate was incubated for 1 hour at 37°C and subsequently washed. Substrates A and B were added, and the plate was incubated for another 15 minutes at 37°C in the dark. The colour development was then stopped with Stop Solution and measured at 450 nm and at 620 nm (correction wavelength). After normalization with correction wavelength, duplicates were averaged, and the average optical density of the blank control was subtracted. A four-parameter logistic standard curve was generated, and the testosterone concentration of each sample was calculated (ng/mL).

### Statistical Analyses

Statistical analyses were performed using the packages survival (version 3.2-13) and lme4 (version 1.1-23) in R (version 3.6.1; https://www.r-project.org/). Latency to Attack and percentage of animals attacking were analyzed using Cox Proportional Hazards Model and Pearson’s Chi-squared test with Yates’ continuity correction. Linear Mixed Effect Models were used to analyze the number of attacks and time spent on aggressive, social and non-social behaviours in the RIT. Linear models were used for the analysis of the locomotor activity in the OF, testosterone levels and 5-HT receptors density. The correlation between testosterone level and the number of attacks in the RIT was tested using the Pearson correlation test. Where applicable, data are presented as mean ± standard error. The level of significance was set at p<0.05.

## RESULTS

### pHyp-DBS behavioural effects

In the DBS group, a significantly lower number of animals attacked the intruders (x-squared=106.67, df=2, p<0.0001) with a significantly higher latency compared to Shams and Controls (β=2.35, exp=10.47, SE=0.26, z=8.88, p<0.0001; Figure 2A). In addition, the number of attacks in each encounter was significantly lower in the DBS group (β=18.13, SE=2.87, df=21, p<0.0001; Figure 2B). DBS-treated animals spent less time on aggressive behaviours (β=68.74, SE=11.99, df=21, p<0.0001) and more time on social behaviours (β=-65.79, SE=14.03, df=21, p<0.001) compared to Shams and Controls. No differences were observed across groups in the time spent on non-social behaviours (β=-3.2520, SE=13.89, df=21, p=0.8). Also similar was the total distance travelled (β=723.9, SE=3997, df=45, p=0.6) and velocity in the OF (β=2.41 SE=13.32, df=45, p=0.6), suggesting that the reduction in aggressive behaviour following pHYP-DBS was not a result of impaired exploratory and locomotor activity. Table 1 presents details of behavioural data.

**Figure 2.**
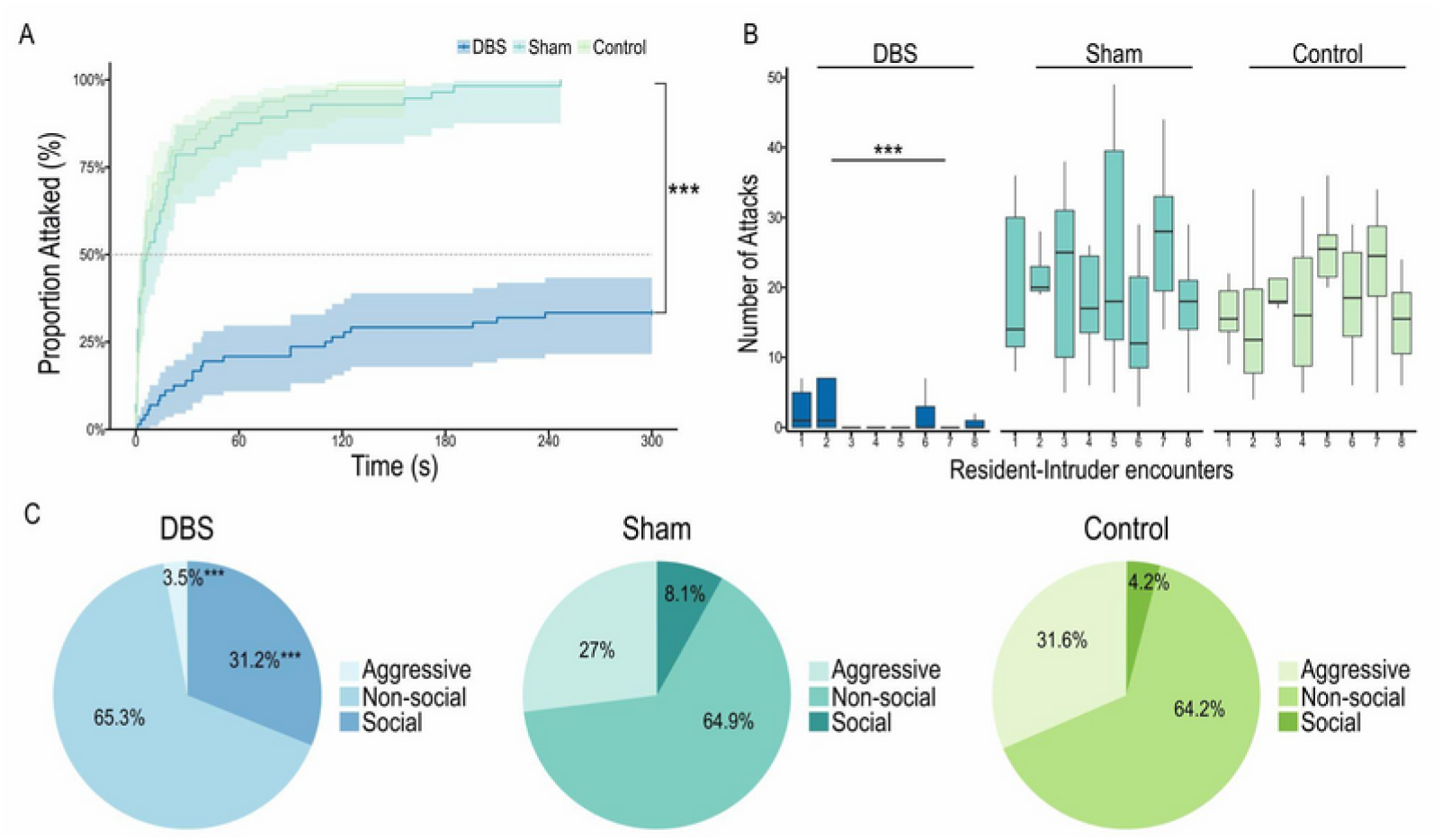
Behavioural results in the resident-intruder test (RIT) in the first experiment. **A**. A smaller proportion of animals attacked with a higher latency in the DBS group compared to Shams and Controls. **B**. Animals in the DBS group attacked fewer times as compared to Sham and Control mice. **C**. Proportion of time spent on aggressive, social, and non-social behaviours. Abbreviations: DBS: Deep Brain Stimulation. In the boxplot, the whiskers indicate the minimum and maximum values; the box indicates the interval from the first to the third quartile, and the horizontal line indicates the median. ***p<0.0001.

**Table 1.** Behavioural response in the Resident Intruder Test and Open Field Test following pHyp-DBS

|  | Variable | DBS | Sham | Control |
| --- | --- | --- | --- | --- |
| RIT | Latency to attack (sec) | 222 ± 118 | 28.2 ± 51.4 | 17.9 ± 31.2 |
|  | Animals attacking (%) | 33 | 100 | 100 |
|  | Number of attacks | 2.4 ± 5.1 | 20.5 ± 11.1 | 18.5 ± 8.9 |
|  | Time in aggressive behaviour (%) | 3.5 | 27 | 31.6 |
|  | Time in social behaviour (%) | 31.2 | 8.1 | 4.2 |
|  | Time in non-social behaviour (%) | 65.3 | 64.9 | 64.2 |
| OF | Total Distance (cm) | 9135 ± 3988 | 9859 ± 4259 | 10464 ± 3765 |
|  | Velocity (cm/sec) | 30.5 ± 13.3 | 32.9 ± 14.2 | 34.9 ± 12.6 |

Abbreviations: DBS: Deep Brain Stimulation, OF: Open Field Test, pHyp: posterior hypothalamus; RIT: Resident-intruder Test. Where applicable, data is presented as Mean ± Standard Error.

### Serotonin receptors 1A and 1B - Orbitofrontal Cortex and Amygdala

DBS-treated animals had an increased 5-HT_1A_ receptor density in the orbitofrontal cortex (β=-4.99, SE=3.19, df=12, p<0.05; Figure 3A) and amygdala (β=-8.02, SE=3.99, df=19, p<0.001; Figure 3B) compared to Sham and Control mice. In contrast, no differences were observed across groups in the density of 5-HT_1B_ receptors in either structure (OFC: DBS: 18.1 ± 6.5; Sham: 16.6 ± 5; Control: 15.2 ± 4; β=-1.51, SE=5.25, df=12, p=0.7; Amygdala: DBS: 14.9 ± 5; Sham: 17.8 ± 3; Control: 17.5 ± 7.6; β=2.88, SE=5.51, df=12, p=0.7).

**Figure 3.**
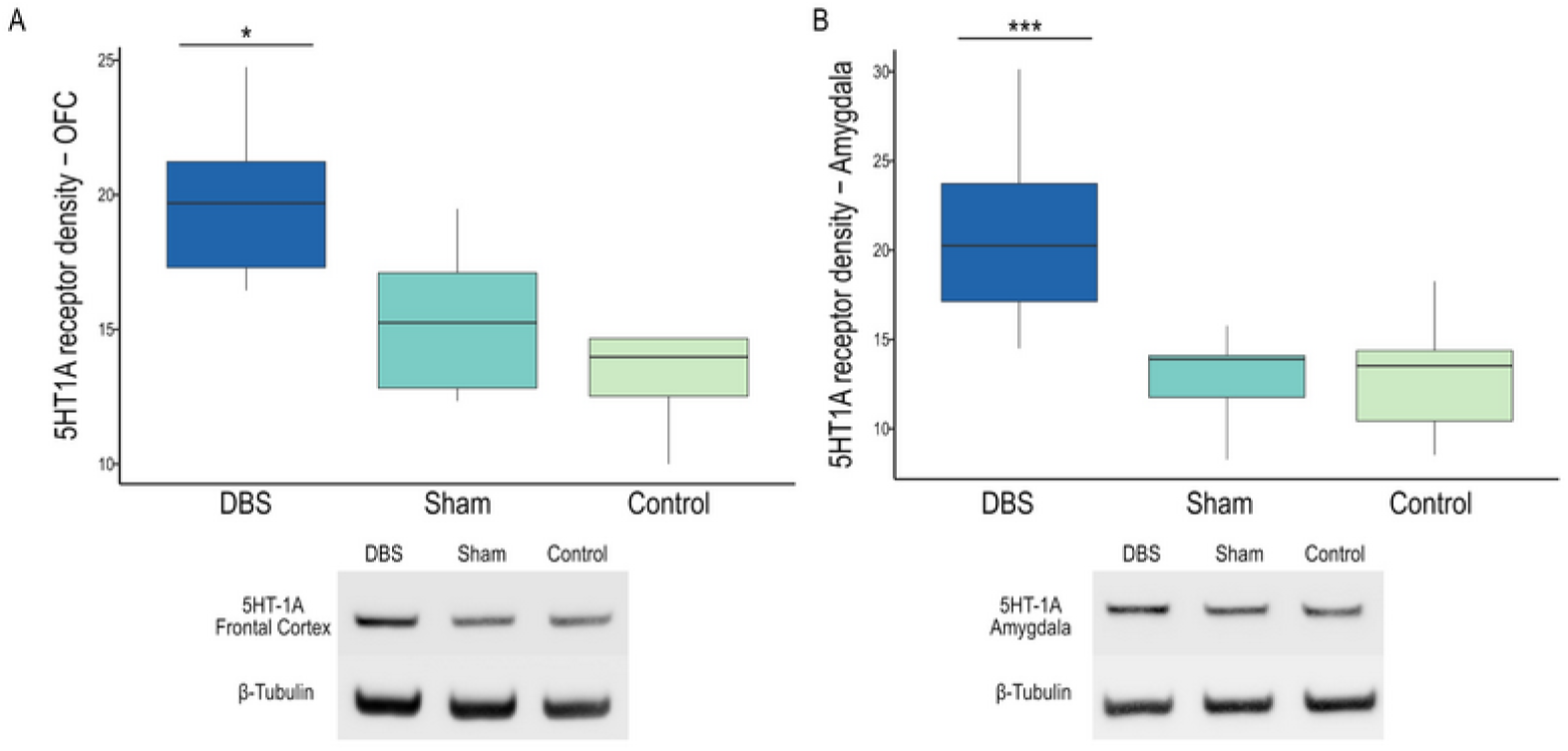
5-HT receptor density. DBS animals showed increased 5-HT1A receptor density in the Orbitofrontal Cortex **(A)** and amygdala **(B)**. In the boxplot, the whiskers indicate the minimum and maximum values; the box indicates the interval from the first to the third quartile, and the horizontal line indicates the median. *p<0.05; ***p<0.001.

### Plasmatic testosterone

Plasmatic testosterone levels were measured in all animals at the last time point. Animals in the DBS groups showed significantly lower levels of testosterone as compared to Shams and Controls (DBS: 12.1 ± 8.5, Sham: 32.7 ± 10, Control: 20.5 ± 5.4; β=12.065, SE=8.111, df=21, p<0.001; Figure 4A). A positive correlation was found between testosterone levels and the number of attacks in the RIT (t=4.39, df=22, p<0.001, R=0.68; Figure 4B).

**Figure 4.**
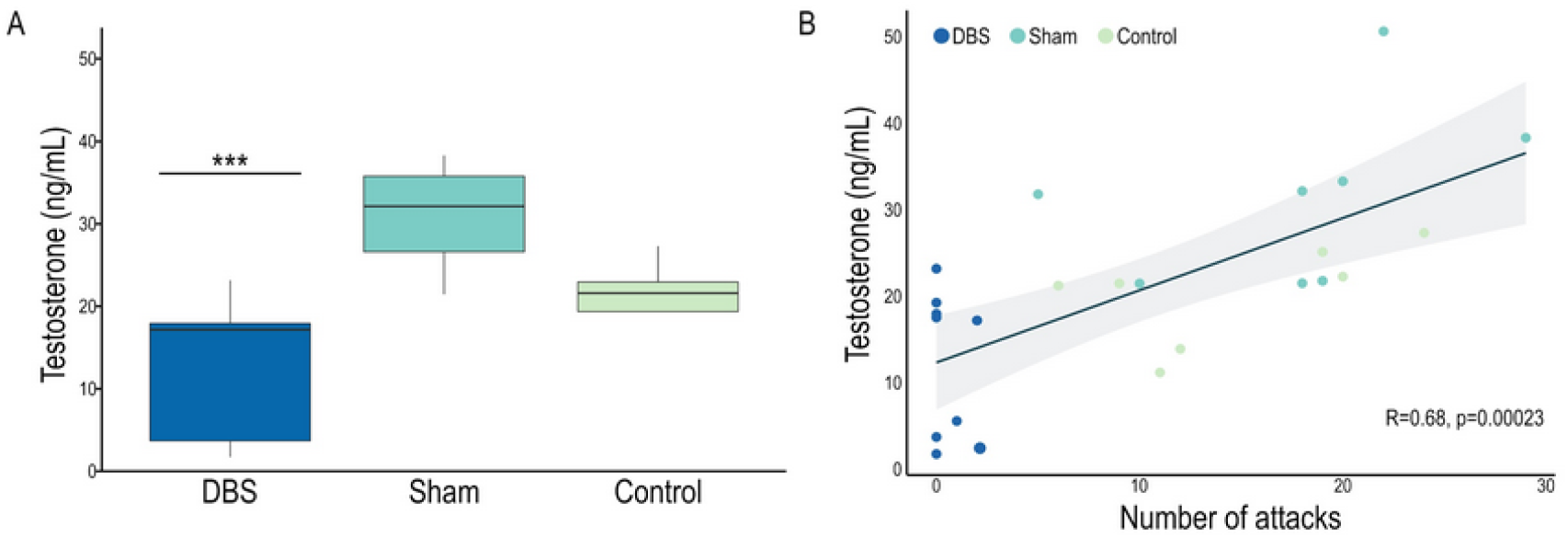
Plasmatic testosterone. A. Plasmatic testosterone levels. B. Positive correlation between the number of attacks and plasma testosterone levels. In the boxplot, the whiskers indicate the minimum and maximum values; the box indicates the interval from the first to the third quartile, and the horizontal line indicates the median. ***p<0.001.

### Pharmacological experiments

Animals attacked the intruder significantly more times when receiving WAY injections before pHyp-DBS (β=-2.75, SE=0.68, df=96, p<0.001) compared to saline-treated controls, regardless of whether the treatment occurred in the first or second half of the experiment (Figure 5).

**Figure 5.**
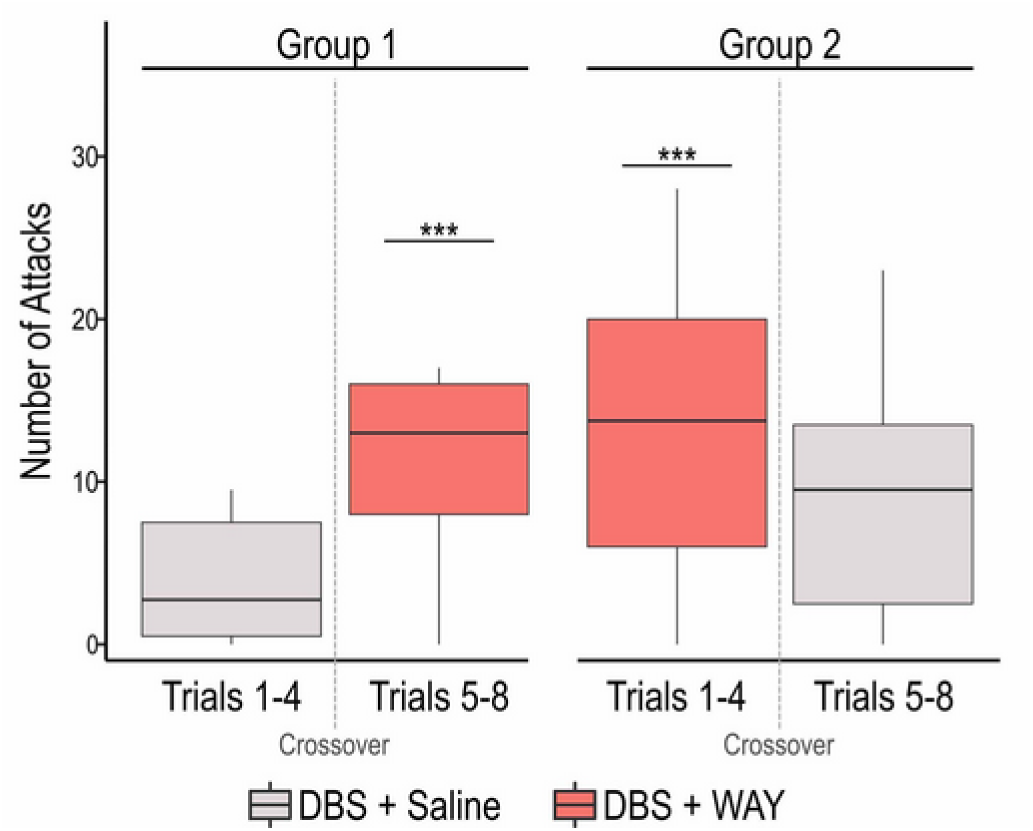
Administration of a 5-HT1A antagonist (WAY-100635 maleate; 2.5 mg/kg SC) before each DBS session blocked the anti-aggressive effect of posterior hypothalamus DBS, regardless of whether it was administered in the first or second half of the experiment. In the boxplot, the whiskers indicate the minimum and maximum values; the box indicates the interval from the first to the third quartile, and the horizontal line indicates the median. ***p<0.001.

The administration of WAY before pHyp-DBS resulted in a higher proportion of animals attacking (x-squared=9.28, df=3, p<0.05), shorter latency to attack (β=0.96, exp=2.61, SE=0.32, z=3.01, p<0.01), more time spent in aggressive behaviours (β=-72.21, SE=22.08, df=84, p<0.01), and less time spent in social (β=41.72, SE=21.13, df=84, p<0.05) and non-social behaviours (β=50.05, SE=24.15, df=84, p<0.05) compared to saline-injected controls (Table 2). In the OF, the administration of WAY before pHyp-DBS resulted in a greater travelled distance (β=50.05, SE=528.1, df=26, p<0.01) and higher velocity (β=-5.77, SE=1.76, df=26, p,0.01) compared animals receiving saline prior to DBS (Table 2).

**Table 2.** Behavioural response in the Resident Intruder Test and Open Field Test following drug injections and pHyp-DBS

| Variable |  | Saline | WAY |
| --- | --- | --- | --- |
| RIT | Latency to attack (sec) | 128±118 | 82.2±111 |
|  | Animals attacking (%) | 73 | 87 |
|  | Number of attacks | 6.7±7.1 | 13.5±11.7 |
|  | Time in aggressive behaviour (%) | 12.7 | 23.5 |
|  | Time in social behaviour (%) | 16.3 | 12.5 |
|  | Time in non-social behaviour (%) | 71 | 64 |
| OF | Total Distance (cm) | 6474 +/- 963 | 8477 +/- 1786 |
|  | Velocity (cm/sec) | 22.5 +/- 3.21 | 28.3 +/- 5.75 |

Abbreviations: DBS: Deep Brain Stimulation, OF: Open Field Test, pHyp: posterior hypothalamus; RIT: Resident-intruder Test. Where applicable, data is presented as Mean ± Standard Error.

## DISCUSSION

Since the 1960s, several preclinical studies have described a critical involvement of the hypothalamus in the expression of aggressive behaviours [17,54,55]. Along with these discoveries, hypothalamic lesions were performed in humans with an average 80% reduction in aggressive behaviour [17]. These surgeries mainly targeted the posterior region of the hypothalamus, in an area known as the “triangle of Sano”, located at the midpoint of the inter-commissural line, the anterior border of the mammillary bodies, and the rostral end of the aqueduct [56]. Hypothalamic lesions, however, have a high potential for severe permanent or transient surgical complications [17,57]. More recently, DBS of the posterior region of the hypothalamus has been investigated for the treatment of aggressive behaviour in patients who exhausted all available medical treatments. Overall, trials in which pHyp-DBS was used to treat aggressive behaviour have shown an average improvement of 91%, with no severe permanent side effects [17,58]. Despite these striking results, the mechanisms responsible for the symptomatic effects of this therapy remain largely unknown.

Translational models of aggressive behaviour offer the possibility of studying neuroanatomical underpinnings and the mechanisms underlying the improvements observed following different treatments [59]. In particular, models of escalated aggressive behaviour aim to mimic excessive, injurious and impulsive aggression that deviates from normal patterns [55,59,60]. These models are based on reduced inhibitory control and are more related to human excessive aggression, such as the one observed in psychiatric patients [59]. One form of modelling escalated aggressive behaviour is via social instigation, where an individual is briefly exposed to a rival before the actual confrontation [59–61]. Social instigation leads to the activation of the limbic system and increases the frequency, intensity and duration of the attacks compared to normal species-typical aggressiveness [59–61].

Here we investigated some of the potential mechanisms by which pHyp-DBS induces its anti-aggressive effects. In the first experiment, we delivered pHyp-DBS before behavioural testing and observed that this treatment was effective in preventing the expression of escalated aggressive behaviour since the first session. In the second experiment, half of the animals received the 5-HT_1A_ antagonist WAY before the DBS session. This drug blocked the anti-aggressive effect of pHyp-DBS and resulted in the expression of escalated aggressive behaviour. After four sessions of WAY administration, animals presenting escalated aggressive behaviour crossed over to the saline arm. In these animals, pHyp-DBS was successful in reducing aggression behaviour, suggesting a treatment effect of stimulation in animals that were already aggressive. These results indicate that pHyp-DBS prevents the development of aggressive behaviour and reduce escalated aggressive behaviour in animals already aggressive.

Our study showed a significant reduction of aggressive behaviours in animals receiving pHyp-DBS in a model of escalated aggressive behaviour that correlated with reduced levels of plasmatic testosterone. In addition, pHyp-DBS resulted in an increased density of 5-HT_1A_, but not 5-HT_1B_, receptors in the OFC and amygdala. Finally, treatment with the 5-HT_1A_ antagonist WAY blocked the anti-aggressive effect of pHyp-DBS.

A correlation between testosterone and aggressive behaviour has been demonstrated in several species, including birds [62,63], rodents [64], non-human primates [65], and humans [66–69]. Testosterone production and release are mainly controlled by the hypothalamic-pituitary-gonadal axis. The secretion of gonadotropin-releasing hormone by the hypothalamus to the anterior pituitary gland leads to a release of luteinizing and follicle-stimulating hormones, which will ultimately act on the gonads to stimulate the production of testosterone [70,71]. At a cellular level, testosterone is aromatized, binds to androgen receptors and then into the respective DNA element, activating or silencing transcription of *cis*-linked genes [62,72,73]. In birds, the activity of aromatase enzymes in the posterior hypothalamus correlates with the intensity of aggressive behaviours [62].

Interestingly, there is a strong correlation between testosterone levels and brain 5-HT availability [46,74,75]. While the administration of testosterone reduces the level of 5-HT and its metabolites in the limbic system [44,45], chemical or surgical castration results in an increase in hypothalamic 5-HT neurotransmission [76–78]. Reduction of 5-HT availability is associated with a series of neuropsychiatric symptoms [79–82], including excessive aggressive behaviour [68,83,84]. Treatment with selective 5-HT reuptake inhibitors (SSRIs), a drug that elevates extracellular levels of 5-HT, can reduce aggressive behaviours in both clinical and preclinical settings [85–88]. These findings indicate that increased testosterone levels and reduced 5-HT availability are associated with aggressive behaviour. Although we did not measure 5-HT levels, the changes observed in DBS-treated mice, with a reduction of plasmatic testosterone and an increase in 5-HT_1A_ receptor density in the OFC and amygdala, suggest a possible interaction between these two systems that may contribute to the anti-aggressive effect of pHyp-DBS.

In mammals, there are seven 5-HT receptor families (i.e., 5-HT_1_ to 5-HT_7_), with five receptors for the 5-HT_1_ family (i.e., 5-HT_1A_, 5-HT_1B_, 5-HT_1D_, 5-HT_1E_, 5-HT_1F_), three for the 5-HT_2_ family (i.e., 5-HT_2A_, 5-HT_2B_, 5-HT_2C_) and two receptors for the 5-HT_5_ family (i.e., 5-HT_5A_, 5-HT_5B_). The 5-HT_4_, 5-HT_6_ and 5-HT_7_ families encompass a single receptor [89]. Previous studies have implicated the 5-HT_1_ and 5-HT_2_ families in the neurobiology of aggression [38,90,91], with strong evidence highlighting the involvement of 5-HT_1A_ and 5-HT_1B_ [92–96]. 5-HT1_B_ receptor knockout mice show increased aggressive behaviour relative to wild-type controls [96], and 5-HT1_B_ agonists reduce aggressive behaviour, an effect that can be effectively attenuated by pre-treatment with 5-HT1_B_ antagonists [61,97,98]. Here we did not observe changes in the density of 5-HT1_B_ receptors in the OFC or amygdala, suggesting the anti-aggressive effects of pHyp-DBS are likely not mediated by this receptor.

Similarly, 5-HT_1A_ agonists reduce aggressive behaviour [98–100], and this effect can be blocked by pre-treatment with selective 5-HT_1A_ antagonists [98]. Interestingly, a reduced density of 5-HT_1A_ receptors has been reported in the OFC and amygdala of animals with a genetic predisposition for aggressive behaviours [92]. In a positron emission tomography (PET) study in humans, lower 5-HT_1A_ binding in the OFC and amygdala was observed in subjects with a lifetime history of aggression [101]. In line with these findings, we observed an increased density of 5-HT_1A_ receptors in both these areas following pHyp-DBS. Moreover, we found, using a pharmacological cross-over assessment, that the administration of the 5-HT_1A_ antagonist WAY effectively blocked the anti-aggressive effect of pHyp-DBS.

Beyond reducing aggression, the administration of 5-HT_1A_ agonists has been previously shown to increase sociability, with no changes in exploratory and non-social behaviours [102], an effect that was similar to the one observed following pHyp-DBS treatment in our study. Animals in the DBS group spent more time on social behaviours (i.e. whole body social sniffing, whole body social grooming), a response that was also blocked by pre-treating the animals with WAY. It is important to note that pHyp-DBS neither reduced the capacity of the resident to interact with the intruder, as the time in social behaviour was higher in this group compared to controls nor did it induce a sedative effect, as no differences were observed between groups in the total distance and velocity in the open field test.

## CONCLUSION

This study showed that pHyp-DBS reduces aggressive behaviour in mice via an intricate mechanism involving a reduction in plasmatic testosterone and increased density of 5-HT_1A_ receptors in the orbitofrontal cortex and amygdala. Further studies are necessary to investigate further the involvement of 5-HT in the mechanisms of action of pHyp-DBS.

## ACKNOWLEDGMENTS

The authors would like to thank the staff of the animal care facility at Sunnybrook Research Institute. This work was supported in part with funds from the Harquail Centre for Neuromodulation, Veteran Affairs Canada (CH) and a Canadian Institutes of Health Research (CIHR) Postdoctoral Fellowship (FVG).

## AUTHORS CONTRIBUTIONS

**Flavia Venetucci Gouveia:** Study design, acquisition, analysis and interpretation of data, drafting and revising the manuscript. **Mustansir Diwan:** Acquisition of data, critically revising the manuscript and providing intellectual content. **Raquel CR Martinez:** Conception and design of the study, critically revising the manuscript and providing intellectual content. **Peter Giacobbe:** Critically revising the manuscript and providing intellectual content. **Nir Lipsman:** Critically revising the manuscript and providing intellectual content. **Clement Hamani:** Conception and design of the study, interpretation of data, drafting and revising the manuscript. **All authors** approved the final version to be published.

## SUPPLEMENTARY FIGURE

**Supplementary Figure 1.**
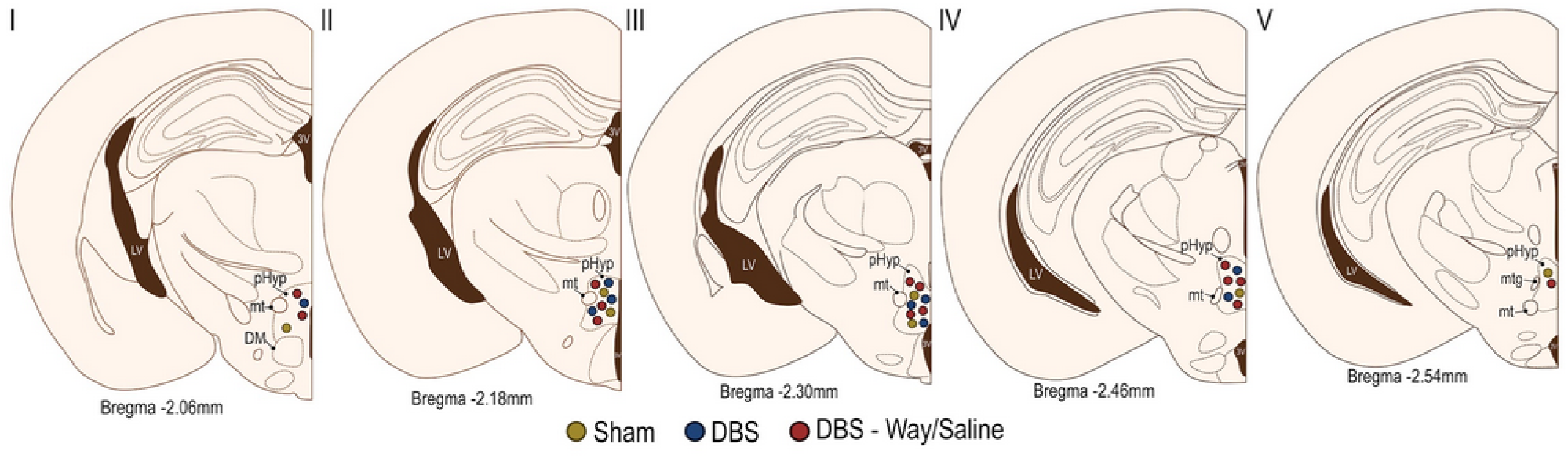
Schematic Illustration of the location of the tips of DBS electrodes. Abbreviation: DBS: Deep Brain Stimulation; WAY: 5-HT1_A_ antagonist WAY-100635 maleate.

